# Temperature effects on reproductive and life-history traits in *Drosophila* (Diptera: Drosophilidae): a systematic map

**DOI:** 10.1101/2025.11.03.686415

**Authors:** Leonel Stazione

## Abstract

Understanding how rising and fluctuating temperatures affect fitness-related traits is critical for predicting the biological consequences of climate change. Ectotherms like *Drosophila* (Diptera: Drosophilidae) are particularly vulnerable due to their physiological dependence of environmental temperature. While many studies have focused on thermal tolerance and survival, sub-lethal temperature effects on reproductive and life-history traits remain understudied despite their ecological and evolutionary relevance. This study presents a systematic review of 288 articles that assessed the temperature effects on six reproductive traits: fertility, fecundity, mating traits, egg hatching, gamete and gonad traits and four life-history traits: longevity, mortality, viability and development time in *Drosophila* species. The results reveal an increase in publications since 1995 and 45 different species employed, with *Drosophila melanogaster* being the dominant model organism. Heat stress was more frequently studied than cold stress, and most experiments involved adult life-stages under constant temperatures. Fecundity and longevity were the most frequently measured traits. Most studies applied constant temperature stress and long-term exposure and few studies examined thermal pre-treatments or artificial selection to evaluate adaptive responses. The data reveals geographical and taxonomic biases, with limited representation from tropical regions and non-model species. This comprehensive dataset underscores the need for broader species coverage, inclusion of variable thermal regimes, and consideration of sex- and stage-specific sensitivities. This study provides a valuable source to guide future studies on thermal adaptation and emphasizes the importance of integrating physiological, behavioral, and ecological data to better predict the response of *Drosophila* and other ectotherms under thermal stress.

## Introduction

Current climate change represents a complex global challenge with great influence on ecosystems and the environment (Adger et al. 2005; Leal Filho et al. 2021; Feliciano et al. 2022). It is having a huge toll on biodiversity, with ectotherms being particularly vulnerable due to the profound effects of temperature on their physiological and ecological processes (e.g. Dillon et al. 2010; Huey et al. 2012).

Climate change is particularly crucial for species with shorter life spans, as they are likely to encounter thermal stressors across a greater proportion of their life cycle (Hoffmann et al. 2013; Kingsolver et al. 2013; Parmesan 2006). In this context, it is essential to study how environmental conditions have shaped biological variation in key traits. Diverse fitness-related traits can be direct phenotypic targets for the thermal adaptation (Hoffmann et al. 2003; Kellermann et al. 2009; Sambucetti and Norry 2015). While assessing survival across thermal gradients remains a standard method for quantifying thermal tolerance (Abarca et al. 2024), there is growing recognition of the importance of sublethal temperature effects -particularly those influencing reproductive and life-history traits-as critical components of thermal sensitivity (Mackay 2002; Melicher et al 2021; Schlesener et al. 2020). *Drosophila* has been widely used as a key model for quantifying the effects of thermal stress on fitness-related traits, revealing adaptive responses to both heat (David et al. 2005; Hoffmann et al. 1997) and cold stress (Chen and Walker 1994; Watson and Hoffmann 1996), with evidence of genetic variability for thermal tolerance. There are several studies about the variation on temperature response, focusing almost exclusively on tolerance under stressful temperatures, while relevant fitness-related traits such as reproductive success and lifespan have received much less attention (Kellermann and Heerwaarden 2019; Walsh et al. 2019).

In natural populations, reproductive fitness is an important target of thermal evolution since thermal stress can reduce organismal activity and lead to mating failure (Miwa et al. 2018). Previous studies have shown that mating success is a relevant component of reproductive fitness (Brooks and Endle 2001; Stazione et al. 2023) and could be a direct target of selection for adaptation to environmental temperature (Dolgin 2006; Sambucetti and Norry 2015). Elevated temperatures can impair both fertility and fecundity, even among individuals that remain fertile (Schou et al. 2021; Walsh et al. 2019). Specifically, previous studies found that fertility and mating success are often correlated under heat-stress environments, whereas this relationship is less evident under benign thermal conditions (Grandela et al. 2024). Cold temperatures can also have detrimental effects on reproductive outcomes (Rinehart et al. 2000; Schou et al. 2021). Cold stress may alter mating behavior and reduce courtship activity or sperm viability, ultimately compromising reproductive output (Anderson et al. 2005; Overgaard and Sørensen 2008). Moreover, exposure to cold may delay reproductive maturation or reduce egg production even in individuals that survive the stress (Watson and Hoffmann 1996; Dolgin et al. 2006). In ectotherms, cold stress may reduce reproductive performance indirectly by lowering activity and growth rates (Pörtner et al. 2000), or directly by impairing gamete function or disrupting embryonic development (David et al. 2005; Watson 2000). Altogether, both heat and cold stress can compromise reproduction and other life-history traits, emphasizing that thermal adaptation involves not only survival under extremes but also the maintenance of fitness-related traits across temperature ranges. The mechanisms underlying thermal effects on reproduction are diverse, reflecting the complex integration of physiological, developmental, and behavioral processes that underlie fertility and fecundity (Walsh et al. 2019). Reported mechanisms include disrupted gonadal development (Delorme and Sewell 2016; McBride et al. 1997), reduced sperm function (Breckels and Neff 2013; Vasudeva et al. 2014), decreased fertilization and fecundity (Hajdu and Hajdu 2022), and reduced resource allocation to gametes or offspring (Berger et al. 2017). While thermal stress can affect reproduction in both sexes, male reproductive traits -particularly sperm production-appear especially susceptible to elevated temperatures (Sales et al. 2018; Schou et al. 2021; Walsh et al. 2019). Conversely, cold exposure can also impair reproductive performance by delaying gametogenesis and mating activity (Watson and Hoffmann 1996; Everman et al. 2018), reducing egg viability, or disrupting embryonic development (David et al. 2005; Rinehart et al. 2000). Importantly, these reproductive processes are often affected earlier during temperature changes, i.e. by less severe conditions than those affecting survival and may have evolutionary and ecological relevance for many populations (Klepsatel et al. 2019; Porcelli et al. 2017).

On the other hand, thermal stress is known to be a determinant of the aging process (Leiser et al. 2011; Gomez et al. 2020). Lifespan is strongly influenced by the metabolic rate of the individual and several studies, in turn, have shown that temperature is one of the most strongly factor modulating metabolic rate in ectothermic animals (Gillooly et al. 2001; Lalouette et al. 2011). Moreover, genotype–environment interactions in life-history traits have been found to affect genetic correlations when different environments are considered (Mackay 2002; Vieira et al. 2000). Leiser et al. (2011) showed that temperature had a significant effect on lifespan of *Drosophila melanogaster* (Meigen 1830) (Diptera: Drosophilidae), showing that lifespan correlates negatively with temperature. Interestingly, another study has shown that inducing mild heat stress may also decrease the aging rate on *D. melanogaster* by stimulating pathways associated with genome stability during hormesis (Sarup et al. 2014). Hormesis in aging positively supports life due to the cellular responses to single or multiple rounds of mild stress (Keil et al. 2015). Additionally, Mołoń et al. (2020) highlighted the fact that there is a clear trend for longer lifespans at lower temperatures in *D. melanogaster*.

Although evolutionary adaptation is typically a slow process, concerns have been raised that many populations will not evolve sufficiently fast to cope with temperature changes (Hoffmann and Sgrò 2011). In this context, phenotypic plasticity in temperature tolerance is often highlighted as an important component of both immediate and long-term responses to fluctuating temperatures (Elmer et al. 2025; Gotthard et al. 2025). Phenotypic plasticity can take many forms and can be induced during development, in adults, and by single or repeated short-or long-term exposures (Rodrigues and Beldade 2020). These plastic responses in ectotherms describe the extent to which single or multiple environmental cues generate changes in their phenotypic responses to novel - and typically stressful-conditions.

Examples of thermal plasticity include acclimation and thermal hardening, particularly under sublethal or moderate temperature stress, where exposure to non-lethal heat or cold conditions can induce protective physiological adjustments. These responses are expected to enhance climate change adaptation by improving survival and physiological performance under heat stress (Sørensen et al. 2019; Simoes et al. 2019). Acclimation for survival is expected to be particularly important during or immediately following extreme thermal events, which can strongly influence the distribution and abundance of insect populations (Everman et al. 2018; MacLean et al. 2019). Further, acclimation and hardening for other physiological and biochemical processes such as metabolic rate can help organisms maintain fitness under fluctuating temperatures (Stazione et al. 2019). In the last decades, several studies have pointed to thermal pre-treatment as a conventional approach to the study of plasticity responses in *Drosophila*, offering insights into how organisms may buffer environmental stress. This is particularly relevant in naturally fluctuating environments, where thermal variability occurs over short (daily) or long (seasonal) time scales, because evolution in such variable contexts is expected to favor high levels of plasticity regardless of whether fluctuations occur within or between generations (Watson and Hoffmann 1996; Bubliy et al. 2013).

Despite the importance outlined above, a large-scale synthesis of the effect of temperature on reproduction and life-history fitness in *Drosophila* is currently lacking. To facilitate such a synthesis, the aim in this study was to build a database of published studies examining the effects of stress temperature on reproduction and life-history traits. A systematic map provides a structured and transparent overview of existing evidence on a given topic, following repeatable ways (James et al. 2016; Haddaway and Pullin 2014). This approach enables the collation, description, and categorization of available studies in a systematic way and can be used for future quantitative analysis or to identify research gaps (James et al. 2016).

To create the map, a systematic search was conducted to identify published studies that statistically examined the temperature effects on both reproductive and life-history traits in *Drosophila*. The search focused on six main types of reproductive trait: fertility, fecundity, mating traits, hatching of eggs and reproductive potential such as gamete and gonad traits, and four main types of life-history traits: longevity, mortality, viability and development time. Overall, a substantial base of evidence was identified, comprising 288 articles that met the inclusion criteria. Based on this literature synthesis, this study examines the effects of climate change on *Drosophila* populations in the context of rising and fluctuating global temperatures, with a particular focus on their reproductive and life-history responses.

## Material and method

### Search method

Online literature searches were performed to investigate how reproductive and life-history traits are influenced primarily by temperature. Therefore, studies retrieved from these searches that did not report temperature were not considered. The searches were performed using the ISI Web of Science Core Collection, Scopus and PubMed platforms considering all available years by adopting PRISMA methodology (Moher et al. 2015). The literature search was conducted up to studies published by September 2025, and unpublished data or grey literatures were excluded.

To be considered eligible for inclusion in the used record, a study had to fulfill the following criteria: 1) Be a peer-reviewed scientific article or book chapter presenting new data, 2) Be conducted on any species of the genus *Drosophila*, 3) Measure at least one of the following reproductive and life-history traits: Gamete traits (sperm or ova number, size, performance, fertilization ability), Gonad traits (testes or ovary size, morphology, developmental stage, function; gametogenesis), Hatching of eggs (proportion of eggs that hatch into larvae, indicating embryonic success), Fertility (the ability to produce offspring), Fecundity (number of eggs or offspring produced), Mating traits (courtship and copulation behavior), Longevity (adult lifespan), Mortality (proportion of individuals dying during a given stage or period), Viability (survival from one developmental stage to the next e.g., egg to adult) and Development time, (duration from egg to adult emergence) 4) Report one of the above traits for at least two different temperatures. 5) Record variation within a species -i.e. multiple individuals are measured for each species-. Review papers, meta-analysis, and opinion articles were excluded. Studies published in any language were considered; however, the search terms were restricted to English.

Both experimental studies, in which individuals were exposed to controlled environmental manipulations, and observational studies, where individuals experienced natural temperature variation, were considered. Studies in which temperature was only indirectly associated with the involved traits -for example, through seasonal changes where date was used as a proxy instead of temperature-were excluded. The search was not restricted to any specific temperature range, and studies involving exposure to both heat and cold conditions were included.

### Screening process

The literature search and study screening process were carried out in successive stages (Figure 1). The initial screening involved 374 articles, and subsequent screenings were conducted based on the following criteria. First, study titles were screened, and irrelevant studies were excluded. Studies that passed the title screening proceeded to the abstract screening phase. Full-text versions of articles passing this phase were then downloaded for a final screening. In this stage, each article was assessed against the inclusion criteria. As a result of this process, 288 studies were identified and included in the final dataset (https://doi.org/10.5281/zenodo.17353764).

**Figure 1.**
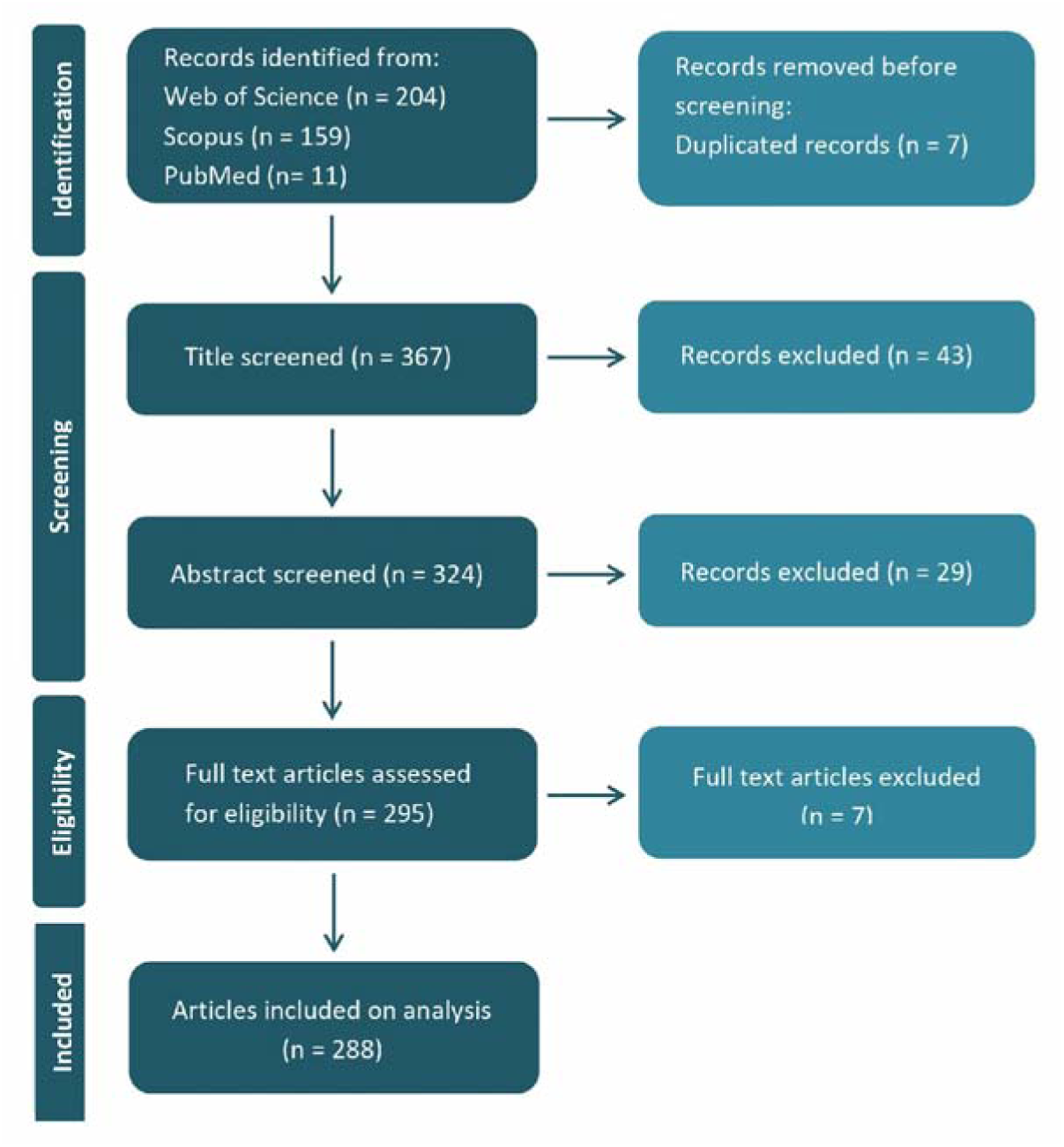
PRISMA diagram flow showing the literature search and study screening process.

### Data extraction and analysis

Following the screening process, the following information was extracted from each of the included articles: Bibliometric information, Species information, Origin of research teams, Reproductive traits, Life-history traits, Sex exposed, Stressor duration, Type of additional stressors, Life-stage, wild or wild-caught status of individuals, Temperature variation, Pre-treatment events, and Artificial selection events. All analyses were performed in the R environment (R version 4.1.2; R Development Core Team 2022).

## Results

A total of 45 species from the *Drosophila* genus were used in all the articles included in the dataset (Figure 2; Table S1). *D. melanogaster* was the most frequently studied species, present in 188 studies, followed by *Drosophila suzukii* (Matsumura 1931) (Diptera: Drosophilidae), present in 29 studies, *Drosophila subobscura* (Collin 1936) (Diptera: Drosophilidae), in 19 studies, and *Drosophila buzzatii* (Patterson & Wheeler 1942) (Diptera: Drosophilidae), in 18 studies (Figure 2, Table S1)

**Figure. 2.**
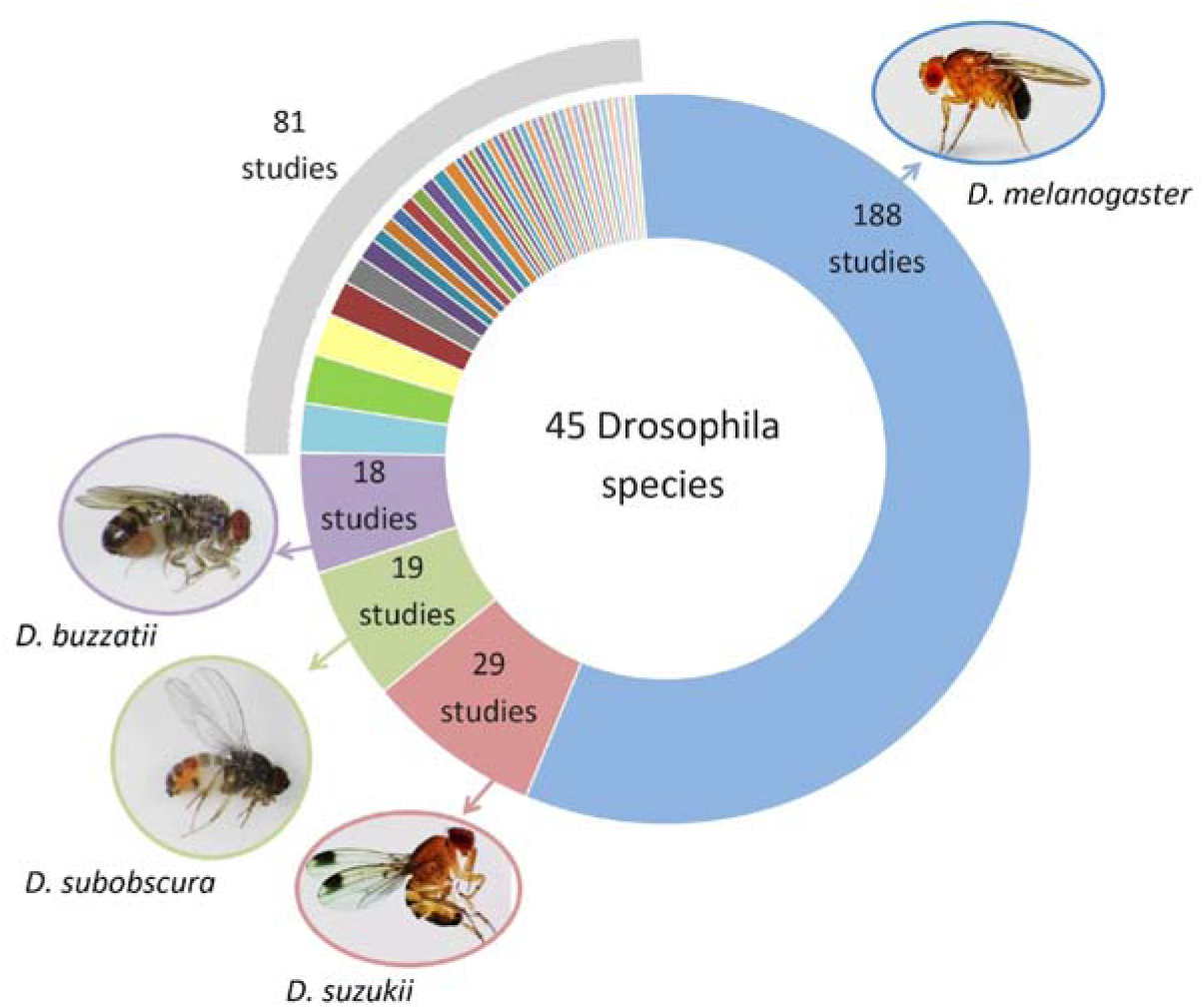
Details of the number of studies that used each of the 45 Drosophila species recorded in the database. The four highlighted species represent the most frequently studied.

The data for the studied traits is summarized in Figure 3. Of the total studies considered, 45.8% (132 studies) recorded measurements of temperature effects on at least one reproductive trait, while life-history traits were measured in 29.9% (86 studies) of the studies. It should be noted that 24.3.7% (70 studies) of the studies were recorded measures of both reproductive and life-history traits (Figure 3A). In the case of reproductive traits, fecundity was the most frequently studied, measured in 42.4% (122 studies) of the studies. Additionally, 19.8% (57 studies) reported temperature effects on mating behavior traits and 16.7% (48 studies) on fertility. In contrast, gonads traits, hatching eggs and gamete traits were the least represented, measured in only 9.7%, 4.9% and 4.9% of the studies respectively (Figure 3B). For the life-history traits, longevity was the most common trait, measured in 31.6% (91 studies) of the studies, followed by mortality measured in 26.7% (77 studies) and viability reported in 15.3% (44 studies). Development time was the least represented, measured in 12.5% (36 studies) of the considered studies (Figure 3B).

**Figure 3.**
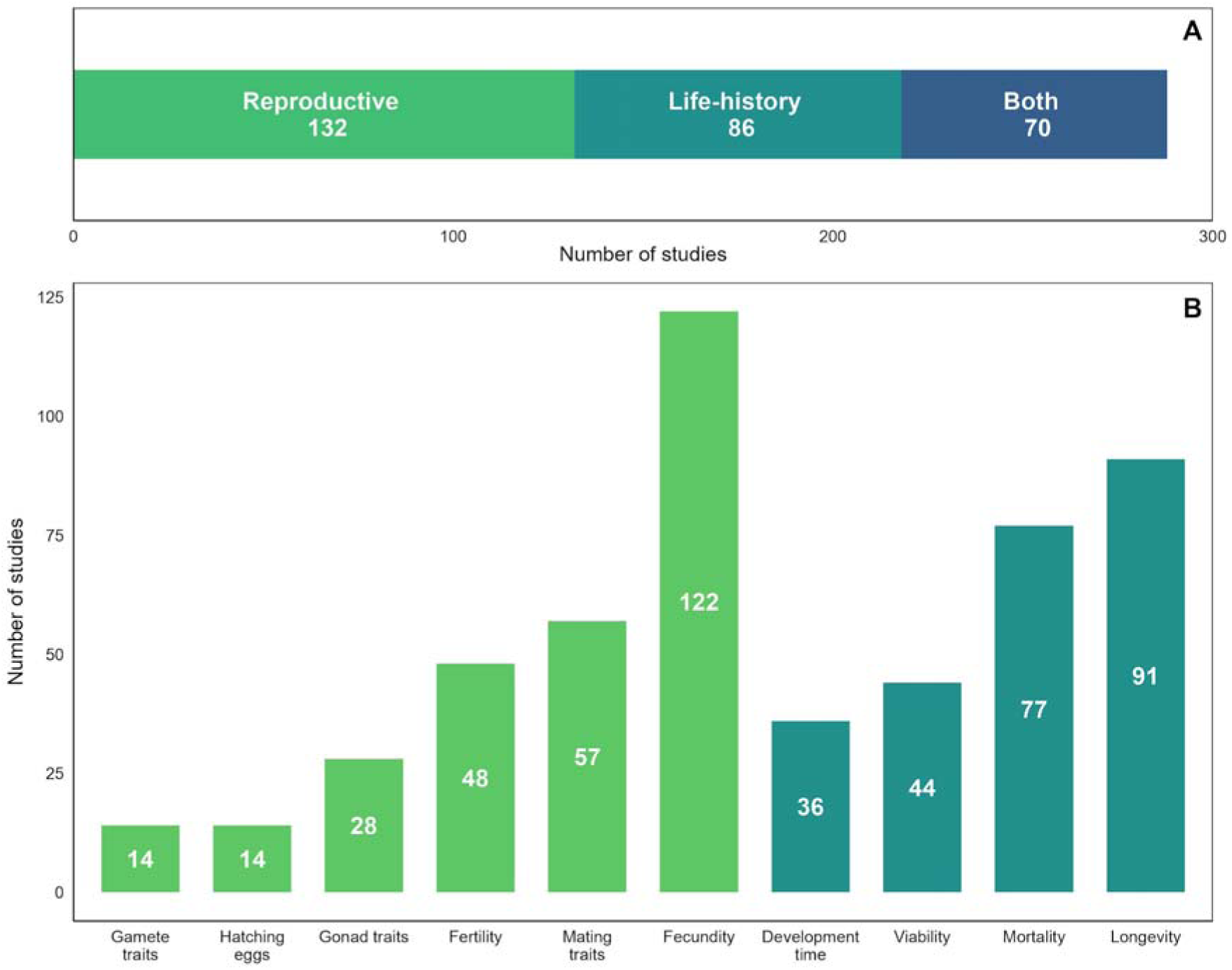
Total number of studies based on the type of traits assessed: reproductive traits only, life-history traits only, and studies that included both traits together (A). Details of the number of studies that included measurements for each specific trait considered in this analysis, see materials and methods (B).

A clear increase in the number of studies published each year after 1995 was observed (Figure 4). The highest number of publications examining thermal stress on reproductive and life-history traits occurred in 2021. The earliest study included in our dataset was published in 1923, and only 35 relevant studies were published before 1995 (Figure 4A). The 288 studies were published across 108 different journals, among which the most common journal categories were “Entomology” and “Ecology and Evolution”, with only seven journals publishing ten or more articles. The most represented journals included Evolution (26 articles), Journal of Evolutionary Biology (17 articles), Journal of Insect Physiology (15 articles), Journal of Thermal Biology (15 articles), and Biogerontology (13 articles; Figure 4B).

**Figure 4.**
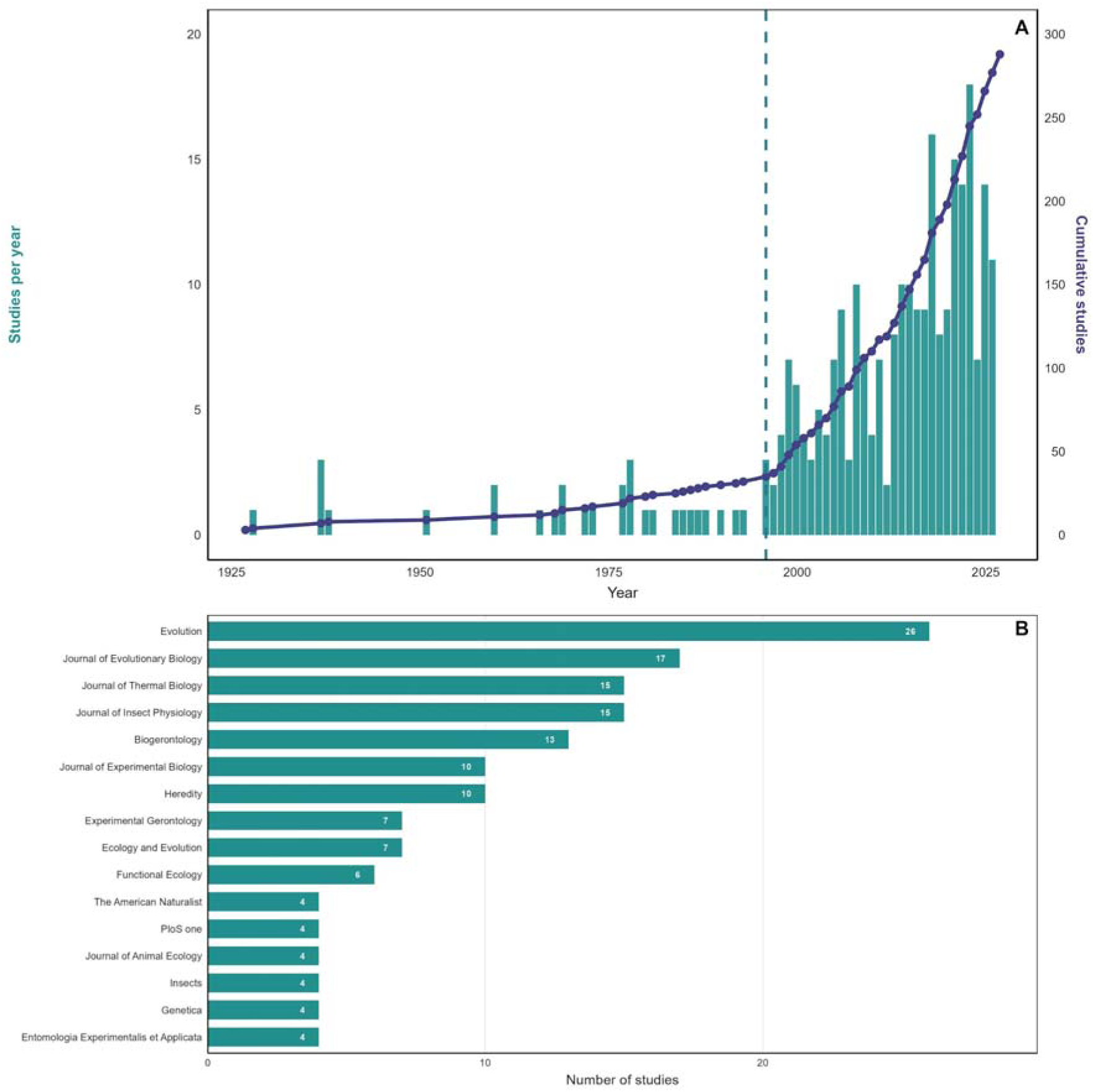
Temporal trends in the number of studies published per year and chronologically accumulated between 1923 and January 2025 (A). And the number of studies recorded by each of the 16 most common journals in the dataset (B).

The methodological characteristics of the experimental studies are summarized in Figure 5. Of the 288 studies, 82.3% (237 studies) exposed both males and females to different temperatures simultaneously, while 8.3% (24 studies) exposed only males, and 6.2% (18 studies) exposed only females. Regarding the duration of thermal stress exposure, 68.4% (197 studies) exposed individuals to thermal treatments for more than five days, 14.2% (41 studies) for one to five days, and 17.4% (50 studies) for less than 24 hours. In terms of life stage, 96.9% (279 studies) applied thermal stress during adult-stage or across several life stages -including adult-, whereas only 3.1% (9 studies) focused exclusively on pre-adult life stages.

**Figure 5.**
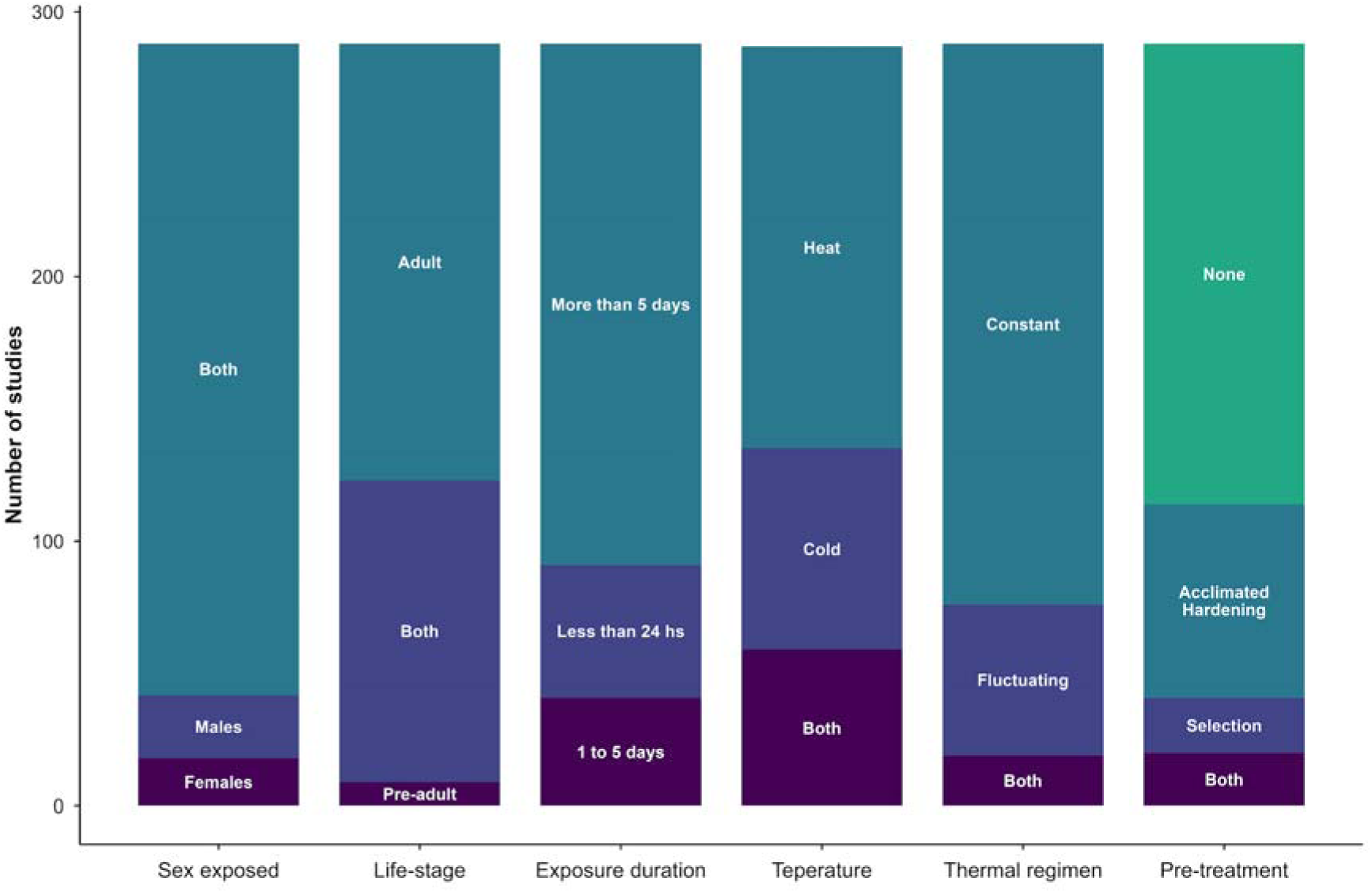
Summary of the methodological criteria considered in the study. The number of studies recorded for each category is shown: sex exposed, life-stage examined, exposure duration, stress temperature, type of thermal regimen, and type of pre-treatment.

In relation to the type of stress applied, 153 studies used only heat stress, 76 used only cold stress, and 59 used both thermal conditions. Concerning treatment regimes, 212 studies used constant temperature treatments, 57 used fluctuating temperatures, and 19 used both constant and fluctuating treatments. Additionally, 41 studies implemented a thermal selection regime on different initial fly populations, and 93 studies applied at least one thermal pre-treatment (e.g., acclimation or hardening) before to thermal stress exposure (Figure 5).

The results indicate that 96 studies used fly populations initially caught from the wild, originating from 35 different countries. The most represented countries were the United States with 20 natural populations, Argentina with 16 populations, Australia with 14 populations, France with 10 populations and Canada with 7 populations (Figure 6A). Additionally, most studies were conducted by research groups based in the United States (68 studies), Denmark (38 studies), the United Kingdom (20 studies), Argentina (20 studies), France (19 studies) and Australia (17 studies; Figure 6B).

**Figure 6.**
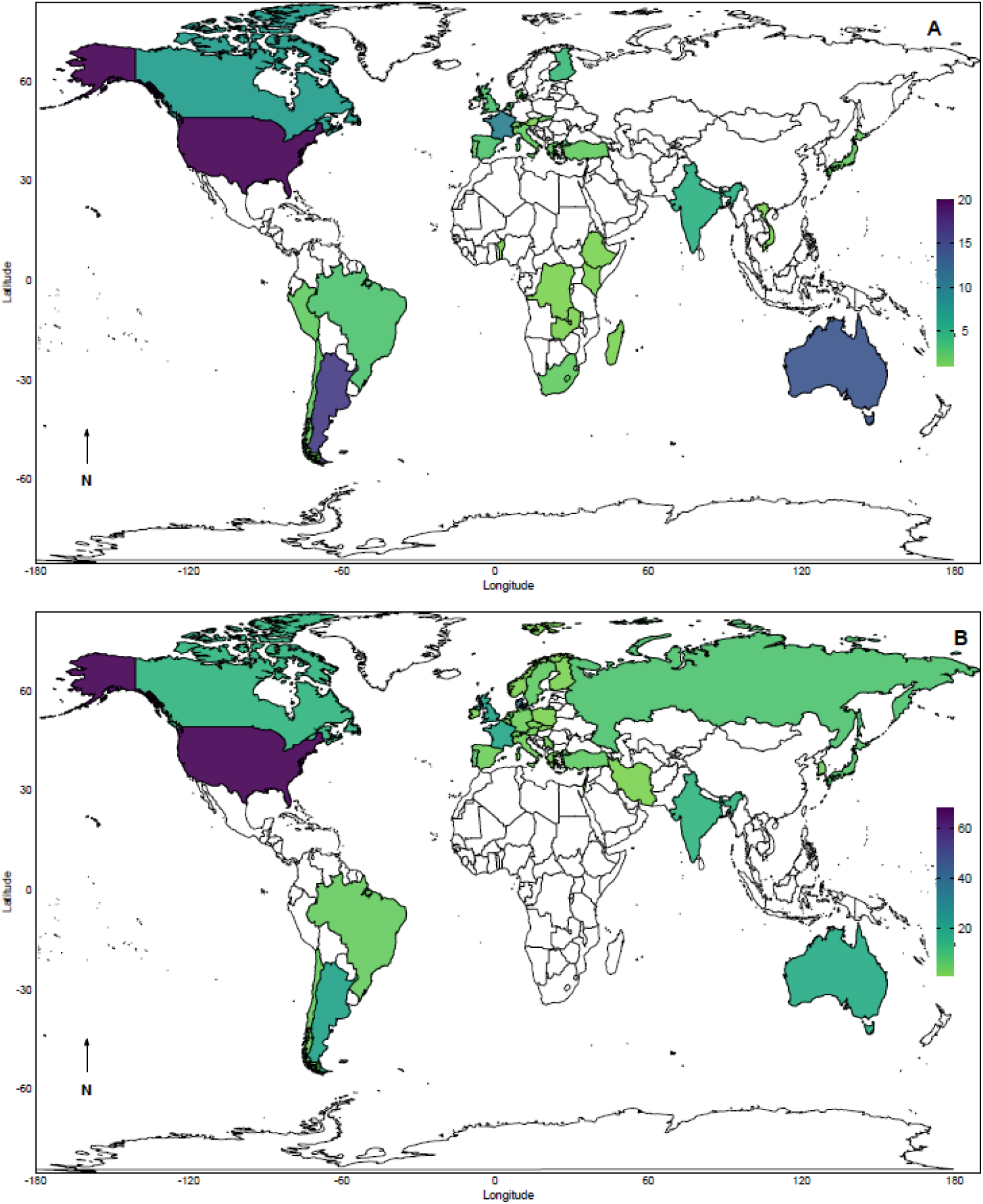
World map showing the number of studies that used fly populations initially caught in the wild in each country (A) and the number of studies conducted in each country, based on the affiliation of the research group (B).

Additionally, a word cloud generated from the titles of all 288 studies revealed that terms such as “resistance,” “temperature,” “reproduction,” “cold,” “selection,” “sterility,” “heat,” “adaptation,” “variation,” “mating,” and “stress” were among the most frequently represented (Figure S1). These terms may serve as useful keywords for searching, identifying and filtering relevant literature.

## Discussion

In this study, a total of 288 relevant articles examining the effects of thermal stress on reproductive and life-history traits in *Drosophila* were identified and analyzed through systematic literature searches. A clear increase in the number of publications has been observed since 1995, with this trend apparently still on the rise. In recent decades the interest in thermal biology has grown considerably, largely driven by the current increase in average global temperatures and the associated increase in thermal fluctuations (Daly et al. 2024; Hunter-Manseau et al. 2025).

Moreover, a higher number of studies focused on thermal stress caused by elevated temperatures, whereas fewer studies examined the effects of cold exposure on fitness-related traits. This discrepancy may reflect the growing concern about understanding the biological impacts of high temperatures, which could be increasingly relevant and necessary in the context of accelerating climate change. This is particularly critical for ectothermic populations living close to their upper thermal limits -maximal temperature allowing individual or population persistence-as they are expected to experience greater vulnerability under fluctuating temperature scenarios (Hoffmann et al. 2013; Nguyen et al. 2011; Payne et al. 2025). These populations must adapt to the changing environmental conditions or migrate to permissive environments to ensure their persistence. Therefore, assessing thermal responses among populations is essential for understanding and predicting the impact of global warming on the distribution of *Drosophila* organisms (Hoffmann and Sgrò 2011; Angilletta 2009). Evaluating the effects of thermal stress on key fitness-related traits, such as reproductive success and lifespan, is fundamental to understanding species response (Klepsatel et al. 2023; Walsh et al. 2021).

More than half of the studies analyzed used *D. melanogaster* as a model organism, likely due to its long-standing history and the extensive body of knowledge associated with this study model. It is known that as model species, *D. melanogaster* has contributed significantly to the understanding of the genetic mechanisms underlying responses to both cold (Sinclair et al. 2007; Bowler and Terblanche 2008; MacMillan and Sinclair 2011; Sørensen et al. 2019) and heat stress (Ashburner and Bonner 1979; Krebs and Loeschcke 1994) through the availability of advanced experimental tools not accessible in non-model species. More recently, genomic and transcriptomic studies have expanded these insights, revealing complex polygenic architectures of thermal adaptation and regulatory responses associated with stress resistance (e.g. Tobler et al. 2014; Porcelli et al. 2015; Zwoinska et al. 2020). Importantly, these approaches have also begun to link molecular mechanisms to variation in reproductive traits, showing that heat-induced fertility loss and changes in gamete function can have a strong genetic basis (Zwoinska et al. 2020; Sales et al. 2018). This reinforces the relevance of D. melanogaster as a key model for integrating physiological, reproductive, and genomic responses to thermal stress.

Nevertheless, other species such as *D. suzukii* and *D. subobscura* were also well represented in the dataset, each with specific scientific relevance. *D. suzukii* particularly is an invasive pest species originally from Eastern Asia, which is now widely distributed across Europe, the Americas, and Africa (Hauser 2011; Cini et al. 2014; Boughdad et al. 2021). There is significant concern over the biological damage it causes, and several studies have explored alternative control strategies to insecticides, such as the Sterile Insect Technique (SIT), for managing *D. suzukii* (Gutierrez-Palomares et al. 2019; Krüger et al. 2019). However, the effectiveness of SIT may be influenced by environmental factors, particularly temperature and humidity (Hamby et al. 2016; Eben et al. 2018; Schlesener et al. 2020). In contrast, *D. subobscura* is recognized as a valuable model species for studying thermal adaptation, due to its broad geographical distribution across the Palearctic region (Rezende et al. 2010). Furthermore, this species exhibits chromosomal inversions that show pronounced variations in frequency that often correspond to temporal and spatial climatic trends (Zivanovic et al. 2023). These latitudinal clines in chromosomal inversion frequencies seem to be responding to the impact of global warming (Rezende et al. 2010). These patterns of natural variation allow studying the influence of historical genetic backgrounds on evolutionary responses (Fragata et al. 2014; Matos et al. 2015). Notably, most studies involving these species were based on sampling from natural populations, which adds greater ecological relevance (Castañeda et al. 2019; Rodríguez-Trelles et al. 2024). Overall, while classical model species like *D. melanogaster* remain invaluable for mechanistic and genomic insights, expanding research to include a broader diversity of *Drosophila* species enhances our capacity to generalize patterns of thermal adaptation and to assess the extent to which plastic and evolutionary responses are shaped by ecological and phylogenetic contexts.

Fecundity was the most studied reproductive trait in the dataset, followed by mating traits and fertility. These three traits are key components of reproductive fitness and therefore essential for understanding thermal adaptation (Mak et al. 2023; Stazione et al. 2023). Although thermal stress has been shown to negatively affect mating ability, several studies indicate that such effects are often temporary, occurring immediately after exposure (Canal Domenech and Fricke 2022).

Fecundity is especially sensitive to both cold and heat stress (Krebs and Loeschcke 1994; Melicher et al. 2021; Rivera-Rincón et al. 2024; Stazione et al. 2017). Interestingly, mild heat stress has been reported to induce hormetic effects -beneficial responses to low-level stress-on male fecundity, comparable to hormetic effects in other fitness-related traits and stress resistance in different *Drosophila* species (Sørensen and Loeschcke 2001; Hercus et al. 2003). In addition, the sterility on individuals carries a very high fitness cost, and may be induced at ecologically-relevant temperatures, suggests that the thermal tolerance of fertility could play a key role in defining current species distributions and extinction risk following warming (e.g. Parratt et al. 2021; van Heerwaarden and Sgrò 2021). It is also important to note that given the short lifespan of the flies a temporary negative effect of mating traits may have crucial importance for lifetime reproductive output. Consistent thermal effects over reproductive traits were found for several recent studies in *D. subobscura*. For example, Simoes et al. (2020) showed that exposure to stressful temperatures during the developmental and adult stage evidenced highly detrimental for reproductive performance in *D. subobscura*. Similarly, Porcelli et al. (2017) showed that individuals from natural populations of *D. subobscura* in both northern and southern Europe exhibited reduced fertility when exposed to high temperatures at development and adult life-stage.

Most studies focused on the effects of long-term exposure (i.e. temperatures applied for more than five days). However, studies examining the effects of temperature on fitness-related traits for short-term temperature were considerably less. Furthermore, most studies examined responses to constant experimental temperatures, with relatively few studies using fluctuating temperature regimes. Considering the high energetic cost of metabolic responses associated with the stress under fluctuating temperatures, the thermal tolerance is a factor of great ecological and evolutionary importance in natural populations, and the correct function of these interacting mechanisms will be a strong determinant of individual survival in stress environments (Manenti et al. 2021). Importantly, environmental conditions in the wild rarely remain constant -even over a single day-thus, studies incorporating some degree of temperature fluctuation could offer a more ecologically realistic perspective (e.g. Rodrigues et al. 2022; van Heerwaarden and Sgrò 2021). Moreover, the results revealed that only a few studies implemented thermal pre-treatments, such as acclimation or hardening, prior to exposing individuals to thermal stress. Previous studies have shown that thermal pre-treatments result particularly important on the individual performance during the thermal stress periods that impact the distribution and abundance of insects (Sisodia and Singh 2006; Arias et al. 2012). In *Drosophila* it is known that there are strong fitness costs and distinct fitness benefits to thermal pre-treatment, depending on the environmental conditions (Everman et al. 2018). For instance, in *D. melanogaster*, previous exposure to sub-lethal temperatures, as in the case of heat-hardening and heat acclimation has been associated to a reduction in mating performance (Stazione et al. 2019), reduction in fertility (Krebs and Loeschcke 1994) and decreasing growth and cell division (Krebs and Feder 1997). The hardening effects as opposed to acclimation are always limited to a short period (e.g., one or few days) in the life of an individual, so a larger effect could be expected for thermal acclimation which results from long-term exposure to a sub-lethal stress (Walsh et al. 2021; Jørgensen et al. 2006).

However, acclimation effects are often limited to specific periods in an individual’s life. Since many ectothermic species live close to their upper thermal limits, the relevance of phenotypic plasticity in meaningfully increasing thermal resistance and mitigating the impact of changing temperatures is currently debated (Barley et al. 2021). Its potential negative effects on other traits could outweigh the benefits, leading to detrimental trade-offs (Kellermann et al. 2012; Sørensen et al. 2016). Therefore, this suggests that a proper evaluation of the beneficial or non-beneficial effects of thermal plasticity in response to environmental changes should consider not only measures of thermotolerance traits but also other fitness-related traits—of significant ecological and evolutionary relevance—that may be indirectly affected.

Only a few studies used artificial thermal selection regimes. However, experimental selection represents a relevant component to study the evolutionary potential of populations in response to environmental challenges (Kawecki et al. 2012). In this way, it is possible to follow the real-time adaptive trajectories of populations and estimating evolutionary rates, tackling how much evolutionary rescue is achievable (e.g. Kinzner et al. 2019; van Heerwaarden and Sgrò 2021). For instance, Mesas et al. (2021, 2023) performed artificial selection for increased heat tolerance in *D. subobscura*, showing that populations can evolve higher knockdown temperatures and thermal resilience within few generations. Interestingly, selection also affected life-history traits, with slow-ramping selected lines displaying higher early fecundity and egg-to-adult viability compared to controls, suggesting pleiotropic effects of thermal adaptation. Also, through experimental thermal selection it is possible to tease apart the effects of plasticity and genetic adaptation during response to stress (Kellermann and van Heerwaarden 2019; Rodrigues and Cogni 2021; Hoffmann and Bridle 2021). For example, selection for cold or heat resistance in *D. melanogaster* has revealed correlated but asymmetric responses between stress types, with evolved populations showing differences in reproductive success, longevity, and development time under stressful conditions (e.g. Gerken et al. 2016; Sørensen and Loeschcke 2001; Singh et al. 2021). Similarly, sexual selection experiments under heat stress in *D. prolongata* showed that thermal conditions can modulate male reproductive success and sperm competition (De Nardo et al. 2025). Several experimental evolution studies on thermal adaptation include genomic analysis, linking molecular and phenotypic variation (Evolve & Reseq). In ectotherms, such studies have revealed a polygenic basis for thermal adaptation (e.g. Tobler et al. 2014; Porcelli et al. 2015). For instance, Tobler et al. (2014) and Porcelli et al. (2015) combined experimental evolution and resequencing in *Drosophila* populations exposed to contrasting thermal environments, revealing rapid allele frequency shifts across hundreds of loci associated with stress response, metabolism, and reproduction. These findings demonstrate that thermal adaptation is highly polygenic and shaped by both direct and correlated selection of fitness-related traits. This study highlights that integrating both experimental selection and genomic approaches could be important to understand the mechanisms driving adaptation to natural environments and the evolutionary potential for thermal adaptation. Overall, experimental selection studies indicate that reproductive and life-history traits can exhibit adaptive responses to thermal stress, although their evolutionary potential is often constrained by trade-offs, asymmetries between stress types, and the complexity of underlying genetic correlations.

Alternatively, studies distinguishing thermal effects across different life history stages will be important to understand the ecological consequences of rising temperatures (MacLean et al. 2019). The effects of temperature on developmental stages have been extensively studied in the literature (e.g. Zhang et al. 2016; Walsh et al. 2019), with direct evidence indicating that both heat and cold tolerance in *Drosophila* are lower during development than in the adult stage (Lockwood et al. 2018). This approach is of great ecological importance by the fact that *Drosophila* species the larval instars are followed by a sessile pupal stage with restricted opportunity to avoid exposure to heat stress, namely through behavioral thermoregulation (Dillon et al. 2009; Rajpurohit and Schmidt 2016). This vulnerability is especially concerning in the context of climate change and the increasing occurrence of heat waves (Kingsolver et al. 2013), which can lead to sudden population declines. For instance, several studies found that thermal stress during development can cause male sterility (Porcelli et al. 2017) because spermatogenesis is more thermally sensitive than oogenesis (David et al. 2005). Therefore, this study underscores the importance of analyzing multiple traits across all life stages to more accurately characterize the thermal limits of natural populations and to identify potential critical periods of sensitivity, particularly during development (Canal Domenech and Fricke 2023).

Another important issue is the differential temperature sensitivity between males and females for the considering traits (Iossa 2019). Most studies in this dataset exposed both sexes to temperature treatments simultaneously. While this approach reflects ecologically realistic conditions, precludes estimates of the specific contributions of each sex to fertility loss. Although it is known that males and females often show different physiological and behavioral phenotypes, according to several previous studies, sex-specific responses to thermal stress and their impact on other fitness-related traits, remain still unclear. For instance, male reproduction appears to be more sensitive to heat stress than female reproduction in certain species groups (e.g., David et al. 2005). Also, previous studies found that *D. melanogaster* males showed greater adaptation to heat stress compared to females (Folk et al. 2006). These differences in thermal response between sexes may be attributed to sex-specific expression patterns of required genes in stress response (Moskalev et al. 2011, Tower et al. 2019). In *Drosophila*, sex determination pathways seem to regulate sex-specific patterns in stress adaptation, where females have been described to preferentially require more genes for stress response than males (Moskalev et al. 2011, 2019). The results highlight that future studies could be relevant to assess thermal effects on males and females separately to better understand the underlying sex-specific mechanisms of thermal sensitivity.

More than one-third of the studies in the dataset examined temperature effects in *Drosophila* populations originally collected from the wild. These populations were sampled across all continents, except for Antarctica. Sampling of natural populations was particularly higher in the mid-latitudes of the Northern Hemisphere, especially in North America, Europe, and Asia. However, high sampling rates were also recorded in Argentina and Australia. Further, most of the articles included in this analysis belong to research groups from North America, Europe, and Asia, showing the same tendency found for population samplings. This pattern reflects a broader trend commonly observed in global collection of population data (e.g., de los Ríos et al. 2018; White et al. 2021). While this distribution may suggest disparities in publication rates among countries and regional scientific systems (Salager-Meyer 2008), it might also result from biases in our search methods, which did not include non-English keywords and did not consider grey literature. Whatever the reason, this biased coverage potentially limits our ability to predict global change impacts (White et al. 2021). It also would suggest that the tropics regions -mainly from Africa and South America-are under-represented in the sample, which is especially problematic given that these regions are currently experiencing the largest temperature anomalies due to climate change (Buckley and Huey 2016). Moreover, tropical ectotherms are particularly vulnerable to future warming, as they already live close to their upper thermal limits (e.g., Bennett et al. 2021; Duffy et al. 2022; Johansson et al. 2020). Overall, this synthesis highlights key gaps in our understanding of thermal effects on reproductive and life-history traits in *Drosophila*. Future integrative studies across sexes, life stages, species, and geographic contexts will be crucial to predict adaptive responses and species persistence under climate change.

## Data availability

The datasets are available at https://doi.org/10.5281/zenodo.17353764

