## Supplementary data for "Temperature effects on reproductive and life-history traits in *Drosophila* (Diptera: Drosophilidae): a systematic map"

Leonel Stazione

University of Florence, Department of Agricultural, Food, Environmental and Forest Sciences and Technologies (DAGRI), Firenze, Italy

**Table S1** Details of the number of studies that used each *Drosophila* species included in the dataset.

| **Species** | **Number of studies** |
| --- | --- |
| *Drosophila melanogaster* | 188 |
| *Drosophila suzukii* | 29 |
| *Drosophila subobscura* | 19 |
| *Drosophila buzzatii* | 18 |
| *Drosophila pseudoobscara* | 7 |
| *Drosophila simulans* | 7 |
| *Drosophila americana* | 6 |
| *Drosophila virilis* | 5 |
| *Drosophila bipectinata* | 4 |
| *Drosophila paulistorum* | 3 |
| *Drosophila birchii* | 2 |
| *Drosophila bunnanda* | 2 |
| *Drosophila ezoana* | 2 |
| *Drosophila koepferae* | 3 |
| *Drosophila mojavensis* | 2 |
| *Drosophila montana* | 2 |
| *Drosophila nigrospiracula* | 2 |
| *Drosophila willistoni* | 2 |
| *Drosophila biauraria* | 1 |
| *Drosophila bunnanda* | 1 |
| *Drosophila busckii* | 1 |
| *Drosophila flavomontana* | 1 |
| *Drosophila immigrans* | 1 |
| *Drosophila littoralis* | 1 |
| *Drosophila mauritiana* | 1 |
| *Drosophila mediopunctata* | 1 |
| *Drosophila mettleri* | 1 |
| *Drosophila mulleri* | 1 |
| *Drosophila nasuta* | 1 |
| *Drosophila nepalensis* | 1 |
| *Drosophila novamexicana* | 1 |
| *Drosophila obscura* | 1 |
| *Drosophila prolongata* | 2 |
| *Drosophila pseudoananassae* | 1 |
| *Drosophila repleta* | 2 |
| *Drosophila robusta* | 1 |
| *Drosophila santomea* | 1 |
| *Drosophila serrata* | 1 |
| *Drosophila sproati* | 1 |
| *Drosophila sturtevanti* | 1 |
| *Drosophila subauraria* | 1 |
| *Drosophila sulfurigaster* | 1 |
| *Drosophila triauraria* | 1 |
| *Drosophila unipunctata* | 1 |
| *Drosophila yakuba* | 1 |

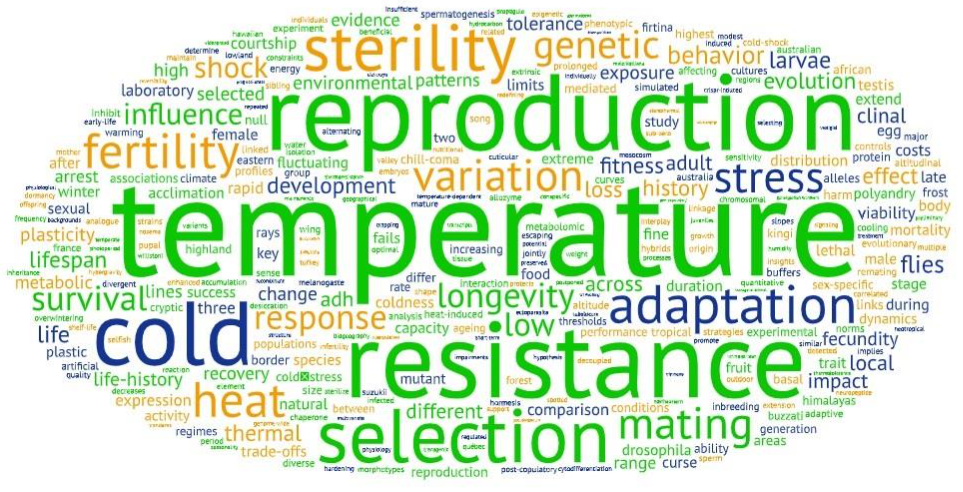

**Figure S1** Visualization of cloud analysis of the single word from the specific titles included in the dataset.
